# Automated Contamination Detection in Single-Cell Sequencing

**DOI:** 10.1101/020859

**Authors:** Markus Lux, Barbara Hammer, Alexander Sczyrba

## Abstract

Novel methods for the sequencing of single-cell DNA offer tremendous opportunities. However, many techniques are still in their infancy and a major obstacle is given by sample contamination with foreign DNA. In this contribution, we present a pipeline that allows for fast, automated detection of contaminated samples by the use of modern machine learning methods. First, a vectorial representation of the genomic data is obtained using oligonucleotide signatures. Using non-linear subspace projections, data is transformed to be suitable for automatic clustering. This allows for the detection of one vs. more genomes (clusters) in a sample. As clustering is an ill-posed problem, the pipeline relies on a thorough choice of all involved methods and parameters. We give an overview of the problem and evaluate techniques suitable for this task.

## I. Introduction

Todays next-generation sequencing technologies enable the analysis of large amounts of genetic information. A number of exciting data sources are given by single-cell sequencing (SCS). Named *Method of the Year 2013* [1], it will be beneficial in many domains of research, most notably medicine and the analysis of disease pathways. Often, on a single cell level, the pathology of complex diseases is very heterogeneous [8]. In different types of cancer, for example, arises the question why certain neighboring cells are malignant while others are not. SCS is able to identify such differences in a high resolution, enabling the analysis of underlying causes and dynamics in great detail, which in turn can be the groundwork for specific treatment.

However, existing SCS technologies are still in their infancy and in order to gain tools with economic relevance, a number of problems need to be resolved. A major potential for development can be seen in DNA isolation. In SCS, samples are taken using patch pipettes or nanotubes. These methods come with the disadvantage that also foreign DNA such as from within the sample (viruses, bacteriophages), or from the laboratory environment can easily be captured [6]. Much effort has been invested in engineering devices for cell isolation and amplification steps that minimize the contamination caused by the surrounding sequencing setup [6]. Still, such measures only decrease the probability for contamination and remaining foreign DNA is detected by tedious manual screening. Additionally, the fast growing amount of data makes this step consuming a lot of time. Therefore, there is a strong need for data analysis techniques that can aid automatic post-sequencing contamination detection. Some species can be detected using supervised methods (i.e. based on sequence similarity to known taxa from databases) and fast classification tools exist [2, 31, 21].

The majority of species is unknown [22] and thus cannot be detected by such methods. Hence, an, taxonomy-free analysis is required [20]. Here, one particularly promising line of research relies on modern techniques from machine learning, specifically clustering techniques based on *k*_*l*_-mer frequencies that already found early applications in metagenomic binning [17]. From the perspective of computational intelligence, contamination detection in SCS is very similar to metagenomic binning. Both metagenomic and SCS samples can be represented as a set of high-dimensional data points using oligonucleotide frequencies. Binning and also contamination detection then correspond to the problem to reliably detect clusters in a high dimensional data space.

In this context, quite a few challenges arise: To circumvent negative side effects in such high dimensional spaces and to enable human expert inspection, it is crucial to use appropriate subspace embeddings to transform the data into an easily visualizable representation, i.e. two or three dimensions. Another challenge consists in the automatic determination of the number of clusters and its cluster validity, a deep and crucial question in the context of clustering [28, 14]. In contrast to metagenomics, in SCS one is concerned with much less genomes in a given sample, significantly reducing the complexity of the problem. Also, contamination detection in SCS corresponds to the problem to discriminate between one or more clusters (genomes). This distinction is important since it heavily reduces the set of applicable clustering methods: The majority of methods for estimating the number of clusters rely on cluster-specific measures such as internal validity measures [18]. Since they are not defined for only one cluster, a distinctive null model for unimodal data is required.

In this contribution, we give and overview on the the-oretical foundations and methodological considerations of an automated contamination detection pipeline. First, we will discuss the suitability of a particular non-linear dimension reduction method. Then, the main focus will be put on evaluating clustering methods and discussing their suitability with respect to different criteria. The outcome of all involved methods depend on a number of parameters. Here, we will suggest strategies for choosing an optimal parameter set. Finally, we show how the inclusion of other, possibly supervised, methods can improve detection accuracy, resulting in a pipeline that can detect contamination with high rate.

## II. Methods

A contamination detection tool will utilize a number of subsequent steps which are outlined in Figure 1. Starting with raw reads from the sequencing process, they are assembled into longer oligonucleotides. Using *k*_*l*_-mer frequencies, a high-dimensional vectorial representation is obtained. After dimensionality reduction, it is the task of evaluating the number of clusters, specifically determining whether there is one or more cluster, resulting in a final decision for contamination. In the following, we will briefly describe the involved methods for frequency computation, dimensionality reduction, and clustering.

**Figure 1:**
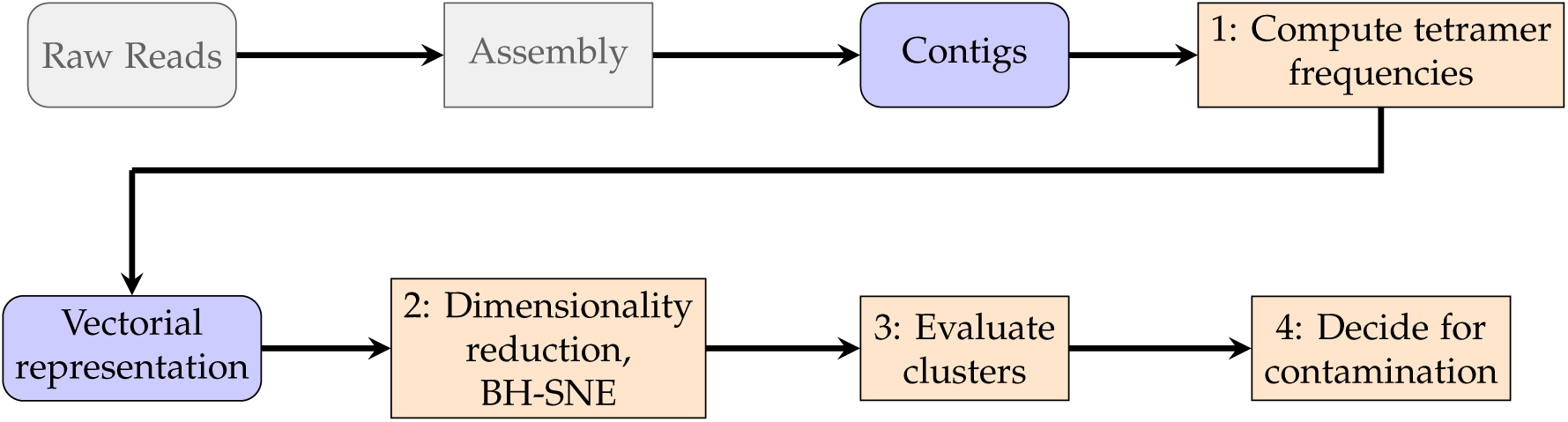
*Contamination detection pipeline.*

### I. Vectorial representation

It is common practice to transfer sequential DNA data into vectors by using signatures of small chunks of DNA [24]. A window of width *w* is fixed and subsequently shifted over the sequence with step Δ*w*. For each shift, the underlying *k*_*l*_-mers frequencies are evaluated. This results in one 4^*k*_*l*_^-dimensional data point per shift, accounting for the 4 nucleotide bases. Exemplary, taking *k*_*l*_ = 4 would result in 256 dimensions, however by accounting for reverse complements, it can be reduced to 136 dimensions.

The choice of window parameters has big influence on the resulting representation. Here, choosing a large window width, capturing genome-specific, rather than gene-specific information will result in less noise [19]. However, a small number of data points is disadvantageous for clustering, such that is has to be taken care to choose *w* not too large. Using a given number of data points, it is possible to estimate the window width and fixed window step accordingly. The default choice of *k*_*l*_ = 4 (tetramer frequencies) usually is robust [19].

### II. Dimensionality Reduction

The analysis of high-dimensional data is often problematic due to the curse of dimensionality [12]. Hence, it is crucial to reduce the dimension while keeping desired properties such as cluster structure.

We employ Barnes-Hut SNE (BH-SNE) [26] as a central method in order to reduce dimensionality in a nonlinear way. It is based on t-distributed stochastic neighborhood embedding [27] which aims to minimize the difference between two distributions of pairwise probabilities in the high and lower dimensional space. Considering *N* data points, high dimensional probabilities are defined as *p*_*ij*_ = (*p*_*i*|*j*_ + *p*_*j*|*i*_) / (2*N*) where

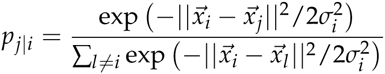

can be interpreted as the probability that 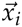 would pick 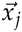 as its neighbor under the assumption that it was picked from a Gaussian distribution centered at *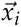*. The parameter *σ*_*i*_ for each data point is automatically determined using a hyper-parameter called *perplexity* that is usually insensitive. Probabilities in ℝ^*d*^ are modeled by

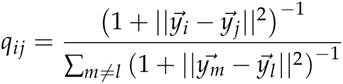

Using the long tailed student-t distribution instead of the Gaussian has the advantage that it allows to avoid the *crowding problem* in low dimensional spaces, leaving more space for distant pairs of points. The Kullback-Leibler divergence between *p*_*ij*_ and *q*_*ij*_ is used to minimize the difference between both probability distributions by numerical optimization.

The original t-SNE method has the major drawback of a quadratic computational runtime and memory complexity, making it unsuitable for larger data sets such as genomes. Barnes-Hut SNE overcomes this deficiency by approximating the similarities between input points, effectively reducing its runtime complexity to linearithmic time while using only linear memory.

BH-SNE has shown to work well for various kinds of data, including genomes [17, 10]. As it puts a particular focus on clusters, in this respect, it is superior to other, possibly linear methods such as PCA. As qualitatively shown in Figure 2, BH-SNE generates clusters which are more compact and separated. A quantitative analysis of the suitability of BH-SNE for clustering DNA sequences can be found in [19]. Clustering algorithms can certainly take advantage of this property.

**Figure 2:**
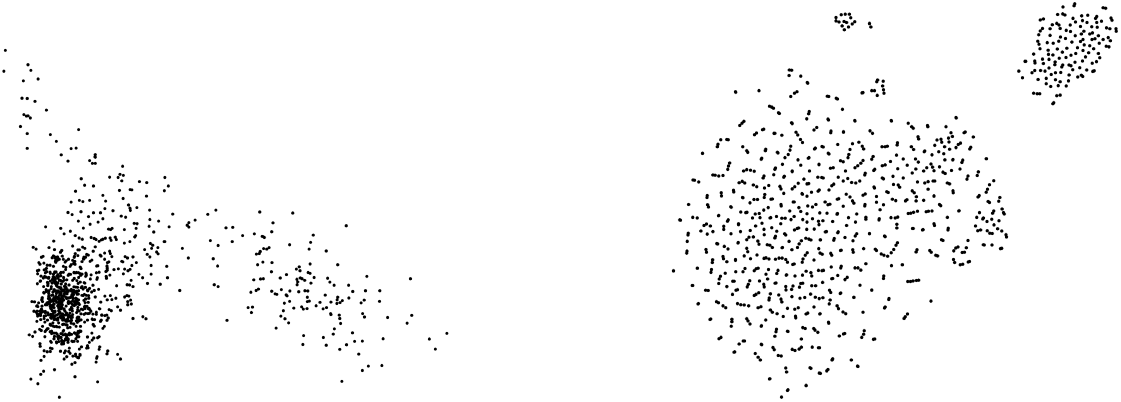
*Comparison of dimension reduction using PCA (left) and BH-SNE (right) of a contaminated single-cell sample.*

### III. Clustering

The goal of clustering is to find a grouping of a given set of data points that is optimal with respect to different objectives. As the notion of a cluster is ill-posed, many different clustering algorithms aim for different objectives [9]. Most techniques depend on a parameter, which is the number of clusters *k*. In some fields, including genome clustering, *k* is unknown and has to be estimated. A special case is given by SCS contamination detection where the determination of the actual number of clusters is secondary. Here, one is not necessarily interested in a specific grouping, rather in the distinction between *k* = 1 (no structure, clean sample) and *k >* 1 (clusters, contaminated sample). For this task, a large subset of *k*-estimation procedures falls out of focus since they operate on cluster-specific characteristics, only defined for *k >* 1. The case *k* = 1 requires an appropriate null model to which the data is compared to in order to be able to detect no structure [25]. In the following, we will review techniques suitable for this task.

We propose a set of differentation criteria for clustering algorithms with respect to the suitability in a contamination detection tool:

- Parameter complexity: As it is difficult to optimize parameters of a given method, even more for researchers from external domains, i.e. biology, the number of parameters should be low. All parameters should be either robust to changes, or easily controllable, possibly through custom hyper-parameters.
- Interpretability of results: Having no absolute truth in clustering, it is desirable for a method to deliver results which are interpretable, possibly including measures of confidence, i.e. p-values.
- Existence of an optimal labeling: The method should be able to provide a grouping for the optimal number of clusters, making it possible to tag all contaminant parts in a given sample.
- Computational complexity: Even single-cell genomes can be very large and samples plenty. For interactive data investigation, methods with long runtime are not desirable.

Given these criteria, we will give an overview of relevant methods and discuss pros and contras. In the following, method parameters are given as *P*_*x*_ where the subscript *x* denotes the actual parameter.

#### III.1 Gap statistic

A very frequently applied method for estimating the number of clusters is the *Gap Statistic* [25]. It considers the difference of the within-cluster dispersion compared to its expected value under a given reference null distribution which is taken to be uniform, aligned at the principal components of the data. *k* is taken according to the largest gap between those measures which are estimated using *P*_*B*_ repetitions. It is found by locating either an elbow or the maximum in the gap curve for a given range 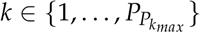. However, the elbow might not be pronounced enough or the maximum is surrounded by a noisy plateau, sometimes resulting in wrong estimations. Even though, the method is statistically well founded, it does not provide any interpretable significance of the result.

#### III.2 Sub-sampling stability

The *Model Explorer* algorithm (ME) and related methods [4, 30] determine *k* by looking at the stability of clusterings for *k ∈* {2,…, *P*_*k*_*max*__} with respect to random sub-sampling of the data. Here, a random subset with fixed ratio *P*_*r*_ is drawn from the data. The number of clusters is chosen as the largest *k* for which a clustering is stable. Here, *stable* is defined as the average similarity (over *P*_*B*_ repetitions) between sub-samples being above a fixed threshold *P*_*t*_0__. If no given clustering is stable, *k* = 1 is assumed. However, this threshold is arbitrary and often depends on the data. Additionally, such methods tend to find stable solutions even on random data [32].

#### III.3 Model Order Selection by Randomized Maps

*Model Order Selection by Randomized Maps* (MOSRAM) [5] can be seen as a variation of the previous Model Explorer algorithm. Instead of sub-sampling, it uses *P*_*B*_ random projections of the data and introduces an additional statistical test for the difference of *k*-clusterings. A set of significant cluster numbers is selected according to p-values of the test using a significance level *P*_*α*_. The method still includes the same, fixed similarity threshold *P*_*t*_0__ as the basis of the test statistic. The random projections take an additional parameter *P*_*€*_ that determines the target dimension. However, in this context, it does not make sense to apply random projections to data which already has been reduced in dimension, however it is worth to note, that the statistical test can also be applied to the Model Explorer algorithm.

#### III.4 Ensemble k-means clustering

The *Multi-K* algorithm [16] randomly samples *k*_*i*_ from a given distribution such as uniformly from *k*_*i*_ ∈ *P*_*K*_ = {1,…, *P*_*k*_*max*__}. It then applies the k-means algorithm *P*_*B*_ times using different *k*_*i*_ in order to build a graph *G* in which edge weights between points of the same cluster are increased in every iteration. In the following, all edge weights are decreased *P*_*B*_ times, in each iteration counting the number of connected components of *G*. The optimal number of clusters is taken as the *k* occurring most often throughout all iterations.

#### III.5 Prediction-based resampling

In *Clest* [7], using a fixed ratio *P*_*r*_, the data is split into training set *L*_*b*_ and test set *T*_*b*_ multiple times *b ∈* {1,…, *B*}. In each iteration, a linear classifier *P*_*C*_ is trained using *L*_*b*_ and tested on *T*_*b*_. At the same time, *T*_*b*_ is also clustered and both the classification and clustering results are compared using an internal cluster validity index. The same procedure is done for a number *B*_0_ of simulated reference null data sets. For each *k ∈*{2,…, *P*_*k*_*max*__}, original and reference performances are compared, resulting in a set of p-values *p*_*k*_. The number of clusters is estimated as *k* for which *p*_*k*_ is significant (using a level *P*_*α*_) and has the largest statistical power.

#### III.6 Dip-means

Using the *Dip Statistic* [11], *Dip-means* [15] is based on a significance test for multimodality. It starts by considering the whole data set as one cluster and tests against the null hypothesis that the cluster is unimodal. The test is applied to each data points distance distribution w.r.t. to all other points within the cluster. If a certain percentage *P*_*v*_ of points has significant evidence against unimodality (using a significance level *P*_*α*_), the cluster is split into two distinct clusters using a clustering algorithm such as k-means. On each resulting cluster, the procedure is performed recursively until no cluster is multimodal anymore. The parameter *P*_*α*_ in combination with *P*_*v*_ can be used to control the number of false positives, detected by the method.

#### III.7 Number of connected components

Spectral clustering can be used to estimate the number of clusters using the eigengap heuristic [29]. However, this heuristic relies on a pronounced elbow in the distribution of eigenvalues that is not distinct enough in most real world data sets, making its identification difficult. Additionally, the construction of the underlying graph heavily influences the result. As BH-SNE (subsection II) often produces very pronounced clusters, we found that counting the number of connected components (CC) of a *P*_*k*_*n*__-mutual-nearest-neighbor graph is often sufficient. This way, counting the number of eigenvalues *λ* = 1 of the normalized graph Laplacian results in the number of clusters as long as they are compact and separated. Tarjans algorithm can be used to estimate this number much faster than using eigendecomposition.

Most of the presented methods deliver an optimal labeling that corresponds to the optimal number of clusters. Only ME, MOSRAM and Clest do not provide such. It is possible to do a posterior labeling. However, it is in no connection to the estimation method, possibly delivering confusing results.

The runtime complexity of all methods is either quadratic or better. For some, it depends most on the number of reference data sets *P*_*B*_ and the underlying clustering algorithm. It is worth to note, that CC can determine *k* in linear time and using Dip-means, it is possible to detect contamination early in the algorithm by only testing for multimodality on the one cluster containing all data.

## III. Preliminary Results

### I. Methods

Early tests on real single-cell data indicate that most methods work reasonably well. However, we found that stability based methods tend to fail to recognize no structure in the data, i.e. non-contaminated samples. This is due to the fact that even in such data, clusterings can appear stable [32], hence we discard these algorithms (Model Explorer, MOSRAM) as suitable candidates. The Gap Statistic also often fails to discover non-structured data. Here, the gap curve shows spurious elbows, even when there is only one cluster in the data. Also, in Clest, the number of parameters seems to lead to unstable results. Although it does work in the majority of cases, its p-values on which basis the decision for *k* is made, are often very near to the significance level, making it very sensitive. Additionally, included parameters are difficult to tune for end users. In Multi-K, even in the presence of more clusters, *k* = 1 cluster is detected too often, resulting in a number of false negatives. This is due to the fact that, in the underlying graph *G*, positive edge weights between members of different clusters might persist for a long time, favoring a single connected component. Estimating the number of connected components of a *P*_*k*_*n*__-mutual-nearest-neighbor graph gives correct results in the majority of cases. Here, *P*_*k*_*n*__ might be interpreted as the minimum number of data points in a cluster, thus is easily interpretable. However, this does hold only for well separated and compact clusters where the largest distance to a nearest neighbor of the same cluster is smaller than the distance to the nearest point in the most nearby cluster. Overlapping or nearby clusters pose a problem for all of the presented methods and are difficult to distinguish from being one. Here, dip-means is standing out as it is also able to discover also overlapping clusters. As long as the structure is significantly multimodal, it is able to detect such.

All of the presented methods work fairly well for estimating the number of clusters for *k >* 1. However, only two methods, counting the number of connected components and dip-means, stand out in their ability to properly differentiate between *k* = 1 and *k >* 1 without too many false detections. Their parameters are few and easily interpretable, making them good candidates for being applied in single-cell contamination detection pipeline. Still, a thorough evaluation of its behavior for this task is required and methods might be modified and combined with other, possibly supervised methods.

### II. Results on annotated data

In order to evaluate the performance of our pipeline, we use simulated single-cell data. The main advantages over using real data from the laboratory is given by a fully correct ground truth and ease of controlling the level of phylogenetic relatedness of included genomes. Here, we expect that quantifying contamination in samples containing remotely related genomes results in a higher detection rate than in samples with very closely related genomes, i.e. from the same species. We employ mdasim [23] to simulate multiple displacement amplification from a given reference sample that is either clean or contaminated. To simulate the subsequent sequencing process, we use ART [13] to generate reads. Finally, contigs are assembled by SPAdes [3]. In our observations we found no difference between using the simulated data and real-world samples, both in data quality and detection rates. Our pipeline reliably detects most contaminated samples with high confidence. Here, easily traceable contamination (i.e. the contaminant is only remotely related) can be detected by counting the number of connected components of a nearest-neighbor graph, as usually, in such cases clusters are very well pronounced. This step can detect most contamination very fast (linear in the number of graph nodes and edges) and further analysis may be skipped.

In uncertain cases (i.e. the contaminant is very closely related), the neighborhood graph might not contain separately connected components anymore. Here, dip-means is employed to test for multimodality of the data. Again, even closely related species can be separated with this approach. Two example are given by Figure 4 and Figure 5. The samples contain two species from the same family and genus respectively. Even though the two graph components are connected by a small bridge, contamination can still be detected by finding a significant multimodality of pairwise distances (*p* = 0). In contrast, Figure 3 depicts a clean sample. Ideally, the neighborhood graph contains only one connected component and the distribution of pairwise distances is unimodal (*p* = 0.12), indicating no contamination.

**Figure 3:**
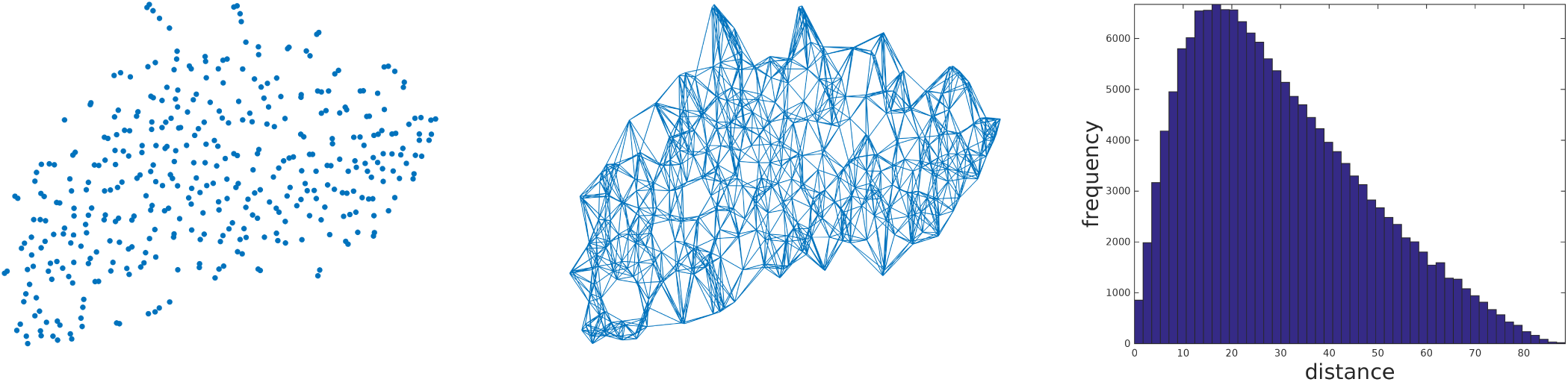
*Cluster analysis of a clean sample. Left: t-SNE representation. Center: 9-nearest-neighbor graph. Right: distribution of pairwise distances.*

**Figure 4:**
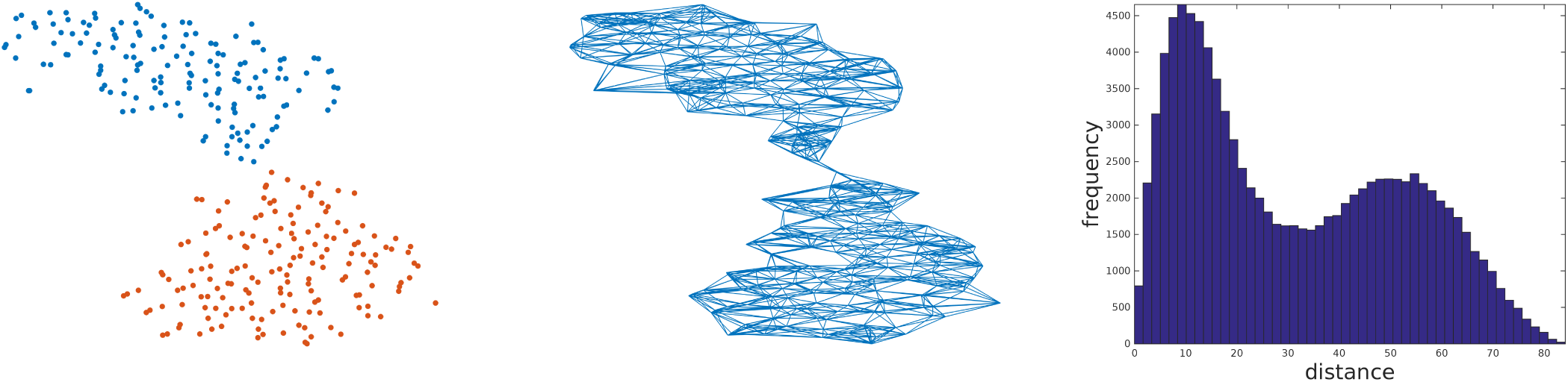
*Cluster analysis of a contaminated sample containing two genomes from the* ***same family*** *(Streptococcaceae). Left: t-SNE representation. Center: 9-nearest-neighbor graph. Right: distribution of pairwise distances.*

**Figure 5:**
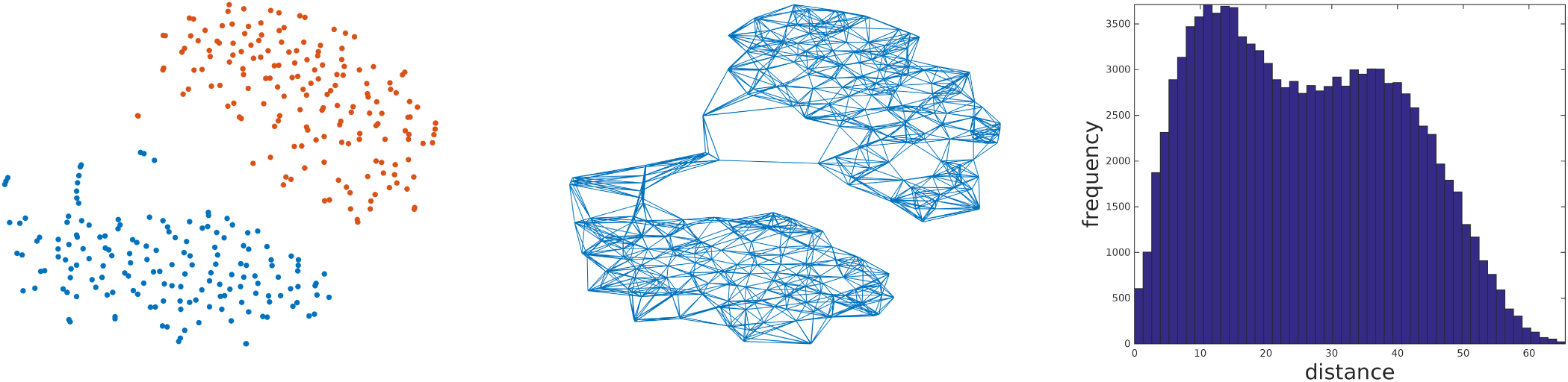
*Cluster analysis of a contaminated sample containing two genomes from the* ***same genus*** *(Streptococcus). Left: t-SNE representation. Center: 9-nearest-neighbor graph. Right: distribution of pairwise distances.*

### III. Summary & ongoing work

Preliminary results are very promising and we plan a more thorough evaluation including many samples with different phylogenetic relatedness. Early observations show that our pipeline can discriminate genomes from the same family and even genus. Because all involved method parameters are determined by the system, our pipeline allows for fully automated batch processing. Still, it allows for interactive inspection by experts and provides *p*-values as confidence. Furthermore, we plan to include various meta data sources such as from the sequencing or assembly process to improve detection accuracy even more.

